# Unveiling the Virulence Mechanism of *Leptosphaeria maculans* in the *Brassica napus* Interaction: The Key Role of Sirodesmin PL in Cell Death Induction

**DOI:** 10.1101/2024.06.15.599173

**Authors:** Marina A. Pombo, Hernan G. Rosli, Santiago Maiale, Candace Elliott, Micaela E. Stieben, Fernando M. Romero, Andrés Garriz, Oscar A. Ruiz, Alexander Idnurm, Franco R. Rossi

## Abstract

*Leptosphaeria maculans* is the causal agent of blackleg disease in *Brassica napus*, leading to substantial yield losses. Sirodesmin PL, the principal toxin produced by *L. maculans*, has been implicated in the infective process in plants. However, the precise molecular and physiological mechanisms governing its effects remain elusive. This study investigates the changes induced by Sirodesmin PL at the transcriptomic, physiological, and morphological levels in *B. napus* cotyledons. Sirodesmin PL treatment upregulates genes associated with plant defense processes, including response to chitin, sulfur compound biosynthesis, toxin metabolism, oxidative stress response, and jasmonic acid/ethylene synthesis and signaling. Validation of these transcriptomic changes is evidenced by several typical defense response processes, such as the accumulation of reactive oxygen species (ROS) and callose deposition. Concomitantly, oxidized Sirodesmin PL induces concentration- and exposure duration-dependent cell death. This cellular death is likely attributed to diminished activity of photosystem II and a reduction in the number of chloroplasts per cell. In agreement, a down-regulation of genes associated with the photosynthesis process is observed following Sirodesmin PL treatment. Thus, it is plausible that *L. maculans* exploits Sirodesmin PL as a virulence factor to instigate cell death in *B. napus* during its necrotrophic stage, favoring the infective process.

**Highlight:** Sirodesmin PL, the principal toxin produced by Leptosphaeria maculans, induces cell death and defense mechanisms in *Brassica napus*, disrupting photosynthesis and facilitating the infective process

## Introduction

*Brassica napus*, also known as canola, is the world’s second-largest oilseed crop, which is highly adaptable and produces large quantities of healthy cooking oil, protein meal, and biofuels as a renewable energy source. As a crucial break crop that performs well in cereal rotations, it is constantly exposed to various microorganisms. In particular, *B. napus* and other *Brassica* species are threatened by Phoma stem canker (blackleg), which is a devastating disease worldwide. This disease is primarily caused by a complex of *Leptosphaeria* species, namely *L. maculans*/*L. biglobosa*, in most producing countries (Zhang and Fernando, 2018). Despite fungicide treatment, these pathogens provoke significant economic losses each season, reducing yield by inhibiting water and nutrient flow through the stem and causing early senescence. In severe cases, the plant stem is severed leading to the death of the plant. Therefore, it is crucial to find effective measures to control the spread of Phoma stem canker in order to maintain the productivity and profitability of *B. napus* cultivation (Fitt *et al*., 2006; Zhang and Fernando, 2018).

Genetic resistance is a highly effective and environmentally friendly method for controlling infections in agricultural crops. Two types of resistance are commonly found in cultivars. Quantitative resistance reduces the severity of symptoms and progression of epidemics over time, but does not prevent infections. This type of resistance is usually controlled by many genes (Huang *et al*., 2009, 2014). On the other hand, qualitative resistance is a dominant resistance that is often effective in preventing pathogens from colonizing plants, and is usually controlled by a single R gene (Rouxel and Balesdent, 2017). In most of these incompatible interactions, the hypersensitive response (HR) is part of the plant’s defense mechanism. During the HR, the plant generates reactive oxygen species (ROS) through oxidative burst, which is associated with the programmed cell death (PCD) of host cells. These responses arrest pathogen growth, particularly biotrophs which rely on living plant tissue (Torres, 2010). However, necrotrophic fungi can subvert this plant defense process in order to obtain nutrients from dead host tissue. For instance, *Botrytis cinerea* takes advantage of ROS and PCD produced after its recognition to proliferate. In this sense, chloroplasts are a primary source of cellular ROS, and any damage to their components can affect the efficiency of photosynthesis and promote the production of defense compounds (Yang *et al*., 2021). Chloroplastic ROS play an important role in *B. cinerea* virulence, as demonstrated by Rossi et al. (2017) using tobacco plants expressing a cyanobacterial flavodoxin (Fld) in chloroplasts. This protein can functionally replace the cognate electron transfer protein ferredoxin (Fd) and significantly suppress ROS build-up in chloroplasts, enhancing plant resistance against *B. cinerea*. Consequently, chloroplasts have emerged as targets for pathogens due to their crucial function in plant immunity. Several microorganisms interfere with chloroplast functioning in an effort to spread throughout the host plants (Yang *et al*., 2021).

Some plant pathogenic fungi are able to produce toxic secondary metabolites as part of a multifaceted strategy to increase their infection and virulence in plants (Möbius and Hertweck, 2009). In some cases, such toxins are able to induce cell death in order to release nutrients or subvert plant metabolism to the advantage of the fungus. Furthermore, toxins derived from pathogens may interact with a wide range of cellular targets, undermine membrane integrity, or alter gene expression, thereby compromising the host plant’s immune system. Some phytotoxins are able to inhibit the activity of plant enzymes, thereby altering the biosynthesis of key metabolites and disrupting normal plant growth and development. Other fungi are able to interfere with plants’ physiology by producing plant hormones, which can also facilitate infection (Möbius and Hertweck, 2009). The diverse array of strategies used by pathogenic fungi to manipulate host plants highlights the complexity of plant-fungal interactions and the importance of understanding the underlying mechanisms involved in these interactions.

The causal agent of Phoma stem canker was previously divided into two isolate categories based on their disease symptoms on *Brassica* species. The presence of sirodesmin PL, a non-host specific toxin, in culture filtrates was used to classify isolates as Tox+ (producing sirodesmin PL and highly virulent) and Tox0 (not producing sirodesmin PL and weakly virulent) (Williams and Fitt, 1999). Later, it was discovered that Tox+ and Tox0 isolates are actually two distinct species being *L. maculans* and *L. biglobosa*, respectively (Shoemaker and Brun, 2001; Fitt *et al*., 2006). Sirodesmin PL is the major component of phytotoxic extracts produced by *L. maculans*. Moreover, most *L. maculans* isolates produce other minor Sirodesmins, including deacetylsirodesmin PL, Sirodesmin J, Sirodesmin K, Sirodesmin H, and phomalirazine (Pedras *et al*., 1990; Pedras and Biesenthal, 1998; Howlett *et al*., 2001). Sirodesmin PL, which causes chlorotic lesions on leaves and possesses antibacterial and antiviral properties (Rouxel *et al*., 1988), belongs to the epipolythiodioxopiperazine (ETP) class of fungal secondary metabolites. These metabolites are characterized by a sulfur-bridged dioxopiperazine ring synthesized from Ser + Tyr. The disulfide bridge enables ETPs to cross-link proteins via cysteine residues and generate reactive oxygen species through redox cycling. These properties are also thought to confer toxicity to Sirodesmin PL (Gardiner *et al*., 2005*b*; Howlett *et al*., 2010). The contribution of Sirodesmin PL to the virulence of *L. maculans* on *B. napus* has been tested by using a Sirodesmin PL-deficient *sirP* mutant (Elliott *et al*., 2007). Inoculations with the *sirP* mutant onto *B. napus* cotyledons cause similar-sized lesions as the wild-type isolate, indicating that Sirodesmin PL is not a virulence factor at this stage of infection. Nevertheless, the mutant causes fewer stem lesions and is only half as effective as the wild-type in colonizing stems. This evidence shows that Sirodesmin PL is an important virulence factor for *L. maculans* that contributes to fungal colonization in *B. napus* stems (Elliott *et al*., 2007; Howlett *et al*., 2010). Curiously, analysis of gene expression indicates the down-regulation of the transcripts encoding the enzymes for sirodesmin biosynthesis in the early biotrophic stage of disease. Enforced expression of these genes renders *L. maculans* unable to cause disease (Urquhart *et al*., 2021).

The mechanism of action of this toxin has not been evaluated, and pertinent studies are necessary to understand the role of Sirodesmin PL as a virulence factor in *B. napus*. In the present study, we evaluate for the first time the morphological, physiological, and transcriptomic alterations of *B. napus* cotyledons treated with Sirodesmin PL. Structural alterations could be observed mainly at the level of the treated cotyledons. On the other hand, the expression of a large number of genes involved in defense mechanisms and detoxification of toxic compounds was induced. In turn, the expression of most genes associated with photosynthetic processes was strongly repressed. This last aspect is correlated with a drop in photosystem II (PSII) activity and the number of chloroplasts. These findings suggest that *L. maculans* employs Sirodesmin PL as a component of its virulence mechanism, leading to cell death in *B. napus*. The research provides a detailed examination of the global gene expression and is accompanied by physiological investigations.

## Material and methods

### Plant material and growth conditions

The experiments were performed using the spring *B. napus* cultivar Westar acquired from IPK Genebank. *B. napus* seeds stored at 4 °C were germinated over wet filter paper and then transplanted to plastic pots filled with a mixture of peat, sand, and perlite (1:1:1), and were irrigated with half-strength Hoagland’s nutrient solution (Hoagland and Arnon, 1950). Plants were grown for 8-10 days in a growth chamber with a 16/8 h photoperiod at 24/20 ± 2 °C and 55/75 ± 5% relative humidity (day/night) and a photon flux density of 200 mmol m^-2^ s^-1^ provided by cool-white fluorescent lighting. At this stage, seedlings consisted only of expanded cotyledons without true leaves.

### Sirodesmin PL purification

Sirodesmin PL was purified and identified as described previously (Callahan and Elliott, 2013). Briefly, Sirodesmin PL was purified from culture filtrate collected from 12-day-old liquid cultures of *L. maculans* grown in V8 juice. The culture filtrate was then extracted twice with ethyl acetate. The organic extracts were dried using a rotary evaporator. The crude extract was resuspended in acetonitrile/H_2_O (1:1) and further mixed. The precipitate was removed by centrifugation at 1,430 × *g* for 5 min followed of filtration through a 0.2 µm Millex-LG PFTE syringe filter to remove any remaining particulate matter. Fractionation was carried out using a Phenomonex Luna 10 × 250 cm 5 µm (ODS) reversed phase semi-preparative column and an Agilent 1100 HPLC system with a UV/Vis detector and fraction collector. Absorbance was monitored at 210 and 240 nm and the fraction eluting corresponding to the elution time of the Sirodesmin PL standard was collected. These fractions were analyzed by ESI-MS/MS as described (Elliott *et al*., 2007).

### Plant treatments

For the bioassays, a 5 mM Sirodesmin PL ethanolic solution stock was prepared by dissolving a known amount of the pure compound in absolute ethanol. Small aliquots of stock solution were kept at −70 °C in a freezer for long periods of time without losing its toxicity. Right before performing each assay, dilutions were made to the desired concentrations in 1% v/v ethanol. Plant treatments were carried out on cotyledons of *B. napus*, using one cotyledon per plant that was detached using a razorblade. Each cotyledon was immediately transferred to a Petri dish containing 0.8% w/v agar-water. To avoid dehydration, the petioles of the cotyledons were carefully immersed in the agar. In all cases, a 10 µL drop of Sirodesmin PL solution or 1% v/v ethanol (control) was applied to a lobe of each cotyledon. Then, a small puncture was made with a hypodermic needle where the drop was placed to facilitate the entry of the toxin. Finally, dishes were sealed and returned to the original growing conditions.

To obtain reduced Sirodesmin PL free of additional thiols, 1 mM Sirodesmin PL was reduced with 5 mM NaBH_4_ for 60 min at room temperature. Native and reduced Sirodesmin PL were diluted at a final concentration of 25 µM (Schrettl *et al*., 2010; Davis *et al*., 2011).

### Trypan blue, dichlorofluorescein diacetate, NBT, DAB and callose staining

Cell viability was analyzed as described previously by Rossi et al. (2011): *B. napus* cotyledons were submerged in staining solution (25 mL of distilled water; 25 mL of lactic acid, 25 mL of glycerol, 25 mL of equilibrated phenol and 1 mL of a 25 mg mL^-1^ solution of trypan blue in water) and incubated for 4 h at 37 °C with gentle agitation (100 rpm). Afterwards, cotyledons were rinsed in water and cleared with chloral hydrate (2.5 g mL^-1^). After rinsing twice with water, samples were equilibrated in 50% v/v glycerol before visualization under a binocular microscope.

Total ROS production was investigated with the redox-sensitive dye 2′,7′-dichlorofluorescein diacetate (DCFDA). For this purpose, detached cotyledons were submerged in 15 µM DCFDA for 30 min after Sirodesmin PL treatments. Green fluorescence was visualized with a fluorescence stereomicroscope (SteREO Discovery.V12, Carl Zeiss) (excitation filter 460 nm; emission filter >515 nm).

3,3-Diaminobenzidine (DAB) staining was performed as described previously in Sašek et al. (2012). Briefly, DAB solution was prepared by dissolving 40 mg of DAB in 200 μL of dimethylformamide and mixing it with 40 mL of water. Detached cotyledons were immersed in the staining solution and vacuum infiltrated. Subsequently, cotyledons were kept at room temperature in a 50 mL tube in darkness until reddish-brown staining appeared. The stained cotyledons were distained in an acetic acid:glycerol:ethanol (1:1:3, v/v/v) solution at 100 °C for 10–20 min. Next, cotyledons were rehydrated in water and visualized with a stereomicroscope.

For detection of O_2_^−^, cotyledons were immersed and infiltrated under vacuum with 1 mg ml^-1^ nitroblue tetrazolium chloride (NBT) staining solution in 10 mM potassium phosphate buffer. After infiltration, cotyledons were kept at room temperature in a 50 mL tube in darkness for 2-3 h. Stained cotyledons were boiled in acetic acid:glycerol:ethanol (1:1:3, v/v/v) solution for 10 min. Samples were then stored in 95% v/v ethanol until visualization with a stereomicroscope. O_2_^−^ was observed as a blue color produced by NBT reduction to formazan.

For callose staining, cotyledons were vacuum infiltrated with an ethanol/lactophenol solution (2:1 [vol/vol]) and incubated at 60°C for 30 min. Samples were then thoroughly rinsed in water and incubated overnight in staining solution (0.01% aniline blue powder in 150 mM K_2_HPO_4_, pH 9.5). Before microscopic analysis, cotyledons were equilibrated in 50% glycerol. Callose deposition was visualized by epifluorescence using excitation filter, 330 to 380 nm; emission filter, 435 to 485 nm (Rossi *et al*., 2011).

### Chlorophyll fluorescence fast-transient analysis

Chlorophyll fluorescence was performed with a portable fluorometer (HandyPEA, Hansatech Instruments, UK) at 16 and 48 HPT following Rossi et al. (2017) with some modifications. Briefly, control and Sirodesmin PL treated cotyledons were pre-darkened for 20 min before analysis using a leaf clip provided by the manufacturer. Subsequently, they were exposed during 1 s to 3000 μmol photons m^−2^ s^-1^ (650 nm peak wavelength) with a dark interval of 500 ms and exposed during 1 s to 3000 μmol photons m^−2^ s^-1^ again and chlorophyll *a* fluorescence was recorded. The fluorescence data were processed by PEA plus software (Hansatech Instruments, UK) to obtain OJIP parameters. A summary of the OJIP parameters used in this study is shown in the Supplementary Table S7.

### Determination of chloroplast number

The number of chloroplasts and their inner structures were evaluated according to Simon et al. (2013). We analyzed three different Sirodesmin-treated and control biological samples composed of three different cotyledons each. An Olympus AX70 light microscope (Olympus, Life and Material Science Europa GmbH, Hamburg, Germany) with a 40× objective lens was used to determine the number of chloroplasts per cell of the palisade cell layer and the spongy parenchyma from semi-thin cross-sections. A total of 100 cells per treatment were examined to calculate the number of sectioned chloroplasts in the cells.

### RNA isolation and transcriptomic analysis

For total RNA isolation, each biological replicate consisted of 10 cotyledon discs from different plants. Each cotyledon disc (10 mm in diameter) consisted of healthy and damaged tissue. Approximately 75% of its surface was healthy and the remaining 25% was damaged. As shown in Supplementary Figure 1, the percentage of damaged tissue was lower 16 hours after Sirodesmin PL treatment. One cotyledon from each replica was used to confirm the performance of the treatment by measuring cell death by trypan blue staining. Total RNA from control and treated samples was extracted after Sirodesmin PL treatment using a Plant Spectrum Total RNA Kit (Sigma), according to the manufacturer’s instructions. Subsequently, samples were treated with the Ambion Turbo RNA-free DNase kit per the manufacturer’s protocol (https://www.thermofisher.com). After DNase treatment, quantity and purity of total RNA was assessed spectrophotometrically in a multi-plate reader Synergy H1 (BioTek). Libraries and RNA sequencing were performed by Macrogen Company (www.macrogen.com - Seoul, Korea) using Illumina technology. Total RNA integrity was checked using an Agilent Technologies 2100 Bioanalyzer with an RNA Integrity Number value greater than or equal to 7. Pair-end sequencing was performed with 101 bp read length. Reads were filtered to remove those corresponding to rRNA contamination, by mapping to *Brassica napus* rRNAs available at the SILVA database (Quast et al., 2013) with Bowtie v1.2.2 (Langmead et al., 2009). Filtered reads were mapped to the *B. napus* genome (Chalhoub *et al*., 2014) using Hisat2 v2.1.0 (Pertea *et al*., 2016) and transcript quantification was performed with the Stringtie program (Pertea *et al*., 2016). Pairwise differential expression analysis between control and Sirodesmin PL-treated samples at both time points used the DESeq2 software v1.22.1 (Love et al., 2014). We considered as expressed, all genes with fragments per kilobase of transcript per million of mapped reads (FPKM) equal or higher than 2. Differentially expressed genes (DEGs) were established with |log_2_(fold-change)| ≥ 1 and q-value ≤ 0.05 as cut-offs.

### Gene ontology (GO) term enrichment analysis

For each DEG, the closest one in *Arabidopsis thaliana* was identified through local BLASTp against ARAPORT11 (Cheng *et al*., 2017). Corresponding *A. thaliana* IDs were used as input for enrichment analysis with AgriGo2 (Tian *et al*., 2017) with default settings. Enriched terms (FDR < 0.05) along with FDR values were processed with GoFigure! Program (Reijnders and Waterhouse, 2021) which summarizes and groups GO terms and generates a graphical output.

## Results

### Sirodesmin PL induces cell death on *B. napus* cotyledons

To determine the impacts of Sirodesmin PL on *B. napus* cotyledons, cv. Westar was selected for study as this is widely used because it does not contain any major genes contributing to qualitative resistance. Symptoms were evaluated by applying 10 µL aliquots of different toxin concentrations on the adaxial surface of cotyledons. Necrotic lesions caused by Sirodesmin PL were observed in a dose-dependent manner 48 hours post treatment (HPT) (Fig. 1A and B>). Tissue death, as shown by trypan blue staining, started as early as 16 hours after 25 μM Sirodesmin PL application on cotyledons and became more pronounced at 48 HPT (Supplementary Fig. S1). Quantification of the necrotic area showed that the toxin caused significantly more cell death than the control treatment (1% v/v ethanol) starting at 10 µM Sirodesmin PL (Fig. 1B). Symptoms of cell death were observed in 100% of the cotyledons treated with concentrations of 10 µM of Sirodesmin PL (data not shown).

**Figure 1.**
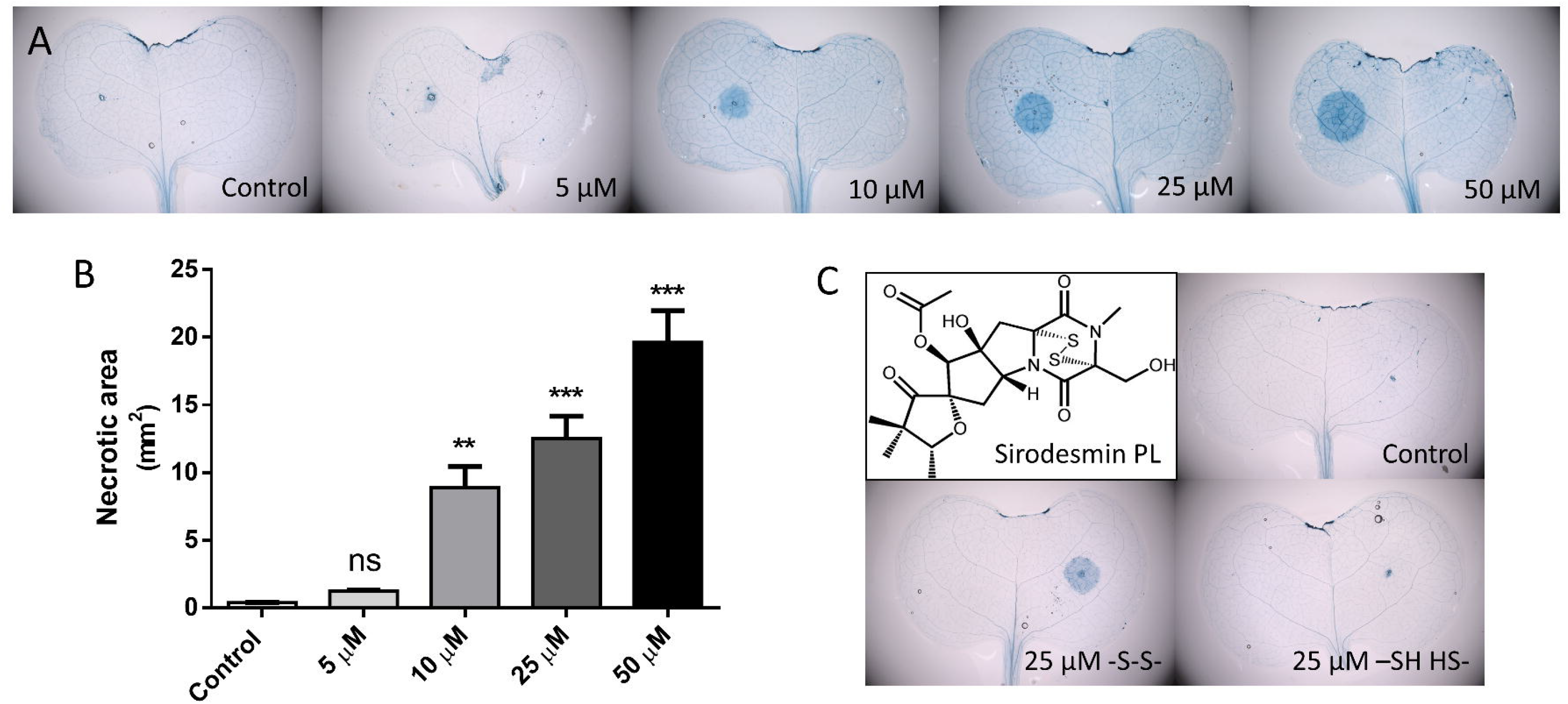
Cell death in *B. napus* cotyledons induced by Sirodesmin PL. Aliquots (10 µl) of Sirodesmin PL solutions in ethanol/water (1:99) were applied to one side of the central vein on the adaxial surface of detached *B. napus* cotyledons. For the control condition, cotyledons were treated with 10 µl aliquots of a solution of ethanol/water (1:99). **A.** Development of necrotic lesions 48 hours after the application of Sirodesmin PL (0, 5, 10, 25, and 50 µM), as evidenced by staining with Trypan Blue and observation under a binocular microscope. **B.** Measurement of lesion size 48 hours after Sirodesmin PL treatment (0, 5, 10, 25, and 50 µM). Results represent the mean of five to seven replicate cotyledons ± standard deviation, and statistically significant differences between different concentrations of Sirodesmin PL and the control condition, as determined by t-test, are indicated as ** and *** (P ≤ 0.01 or 0.001, respectively). **C.** Development of necrotic lesions 48 hours after Sirodesmin PL treatment with 25 µM of oxidized (-S-S-) and reduced (-SH HS-) Sirodesmin PL, as evidenced by staining with Trypan Blue and observation under a binocular microscope.

The native disulfide bond in other ETP toxins, such as gliotoxin, is necessary for both eukaryotic cell uptake and toxicity induction (Schrettl *et al*., 2010). Here, we evaluated the role of oxidation states in Sirodesmin PL’s toxicity on *B. napus* cotyledons. Reduction of Sirodesmin PL was performed using 50 mM NaBH_4_. Treatment with 25 µM of reduced (dithiol form, −SH HS−) Sirodesmin PL did not induce cell death (Fig. 1C), whereas treatment with 25 µM oxidized (disulfide bond, −S-S−) Sirodesmin PL caused similar toxicity as shown previously for 48 HPT (Fig. 1A). As expected, untreated control cotyledons did not show cell death (Fig. 1C). Therefore, the disulfide bond in Sirodesmin PL is necessary to induce cell death in cotyledons of *B. napus*.

### Sirodesmin PL induces global changes in *B. napus* gene expression

RNA-sequencing was used to interrogate the potential changes in gene transcript levels after treatment with Sirodesmin PL. We treated *B. napus* cotyledons with 10 μL of 50 μM of the toxin or 1% (v/v) EtOH (used to dissolve the toxin) for comparison. We took samples at 16 and 48 HPT and performed high-throughput transcriptome sequencing (RNA-seq). We observed low levels of ribosomal contamination, ranging from 0.23% to 1.57%, and a high number of total mapped fragments derived from mRNAs (Supplementary table S1). Fragment mapping ranged between 18,635,301 and 28,437,307 (Supplementary table S1), leading to more than 80% mapping rate. Using Pearson correlation analysis we observed high coefficients between biological replicates belonging to the same treatment (R^2^ > 0.95 in most cases, Supplementary Table S2).

For practical purposes, we considered as expressed all genes with fragments per kilobase of transcript per million of mapped reads (FPKM) equal or higher than 2, in at least one condition. Based on this consideration, we identified 39,258 and 35,791 expressed genes at 16 and 24 HPT, respectively (Supplementary Table S3 and S4). This means that the amount of expressed genes represents approximately 37% of the 101,040 transcripts predicted in *B. napus* annotation v5.

### The expression of thousands of genes are modified in response to Sirodesmin PL

We identified differentially expressed genes (DEGs) by comparing gene transcript levels in mock- and Sirodesmin PL-treated *B. napus* cotyledons considering the expressed genes (FPKM ≥ 2) for each time point after treatment. As a cut off we set |log_2_(fold-change)| ≥ 1 and *q*-value ≤ 0.05. A total of 5,833 DEGs were identified after Sirodesmin PL treatment at 16 HPT, with 3,650 genes upregulated and 2,183 genes being downregulated (Supplementary Table S3). At 48 HPT, from a total of 4,281 DEGs, 2,657 genes were upregulated and 1,624 downregulated (Supplementary table S4). In both time points, we observed more up-than down-regulated genes when the tissue was treated with the toxin. In order to understand in depth the effect of Sirodesmin PL, we analyzed the degree of overlap among DEGs at the two time points studied. As depicted using Venn diagrams, 1,502 upregulated DEGs (31.3%) were shared at both time points (Fig. 2A), while 2,148 and 1,155 genes were specific to 16 and 48 HPT, respectively. In the case of downregulated DEGs, the fraction of overlapping genes was smaller (16.2%) (Fig. 2B), while 1,653 and 1,094 genes were specific to 16 and 48 HPT, respectively. Gene expression values of each subgroup can be found in Supplementary Table S5.

**Figure 2.**
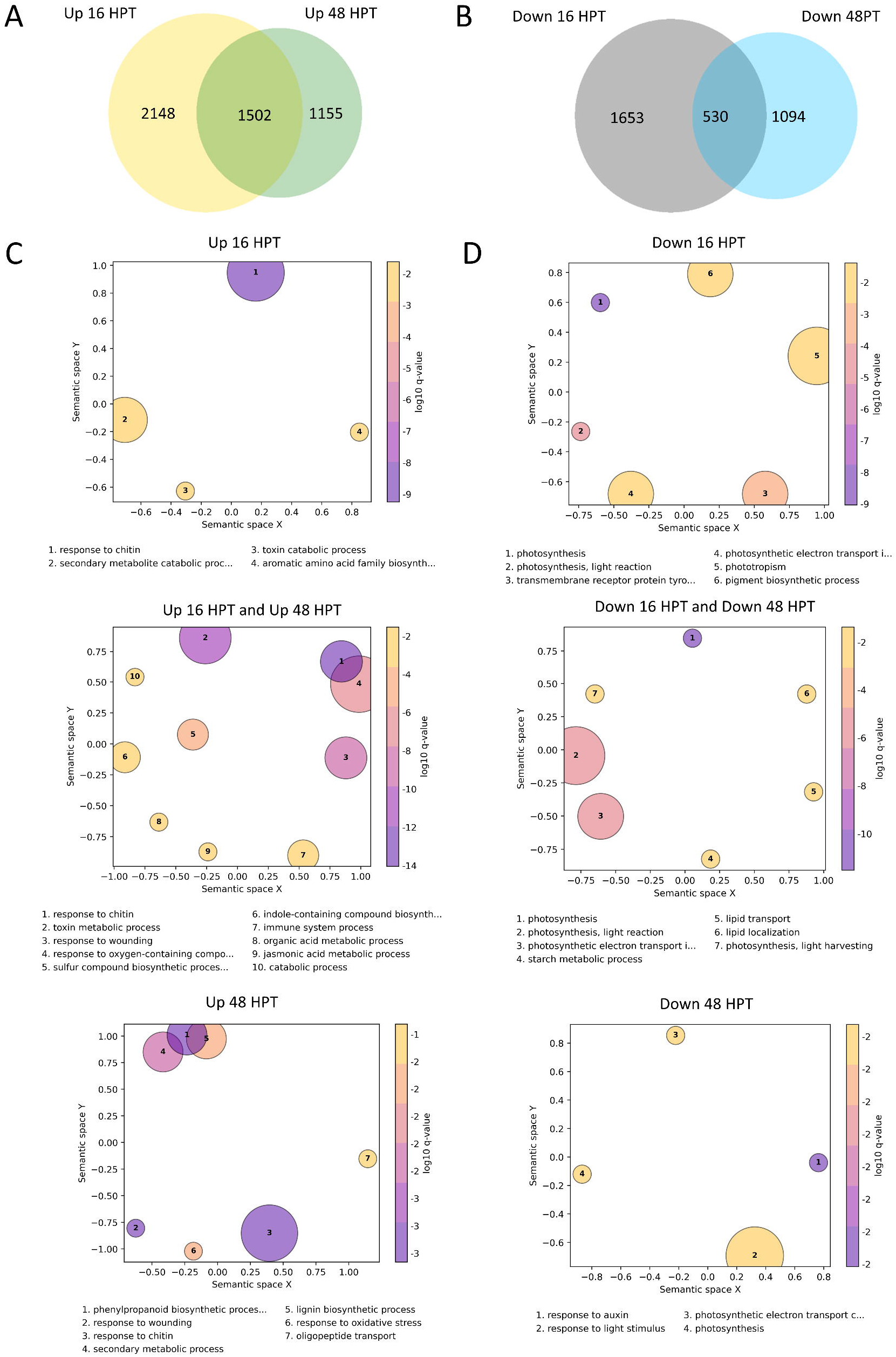
Differentially expressed genes (DEGs) and Gene Ontology (GO) analysis. RNA-seq analysis was performed on *B. napus* cotyledons 16 and 48 hours after the application of 50 µM Sirodesmin PL. A-B. Venn diagram showing the up- (**A**) and down- (**B**) regulated DEGs. After comparing gene expression levels in mock- and Sirodesmin PL-treated B. napus cotyledons, considering the expressed genes (FPKM ≥ 2) at 16 (left) and 48 (right) hours post-treatment, or shared between both times (intersection). GO terms are filtered and grouped by semantic similarity. Bubble size represents the number of genes associated with each term. All GO terms represented in the figure can be found in Supplemental Table S6. **C**. Up-regulated GO terms enriched in cotyledons at 16 (upper panel) and 48 (lower panel) hours post-treatment, and shared at both times (middle panel). **D**. Down-regulated GO terms enriched in cotyledons at 16 (upper panel) and 48 (lower panel) hours post-treatment, and shared at both times (middle panel).

### Gene Ontology analysis support the implication of Sirodesmin PL in plant defense and photosynthesis

The list of DEGs in each subgroup of Venn diagrams (Fig. 2 A and B, Supplementary table 5) was used to perform a Gene Ontology (GO) enrichment analysis. The AgriGO platform (http://systemsbiology.cau.edu.cn/agriGOv2/) was used to classify the up- and down-regulated genes based on their assigned terms. Though all data are available in Supplementary Table S6, we particularly found biological process terms most informative. Statistically enriched terms (FDR < 0.05) from AgriGO output were summarized, grouped and graphically visualized using Go-Figure! software (Reijnders and Waterhouse, 2021). Regarding genes induced by the toxin, enriched terms that can be identified both at 16 and 48 HPT include ‘response to chitin’, ‘sulfur compound biosynthetic process’, ‘immune system process’, ‘toxin catabolic process’, ‘jasmonic acid metabolic process’ and ‘response to oxygen containing compound’, among others. The ‘aromatic amino acid family biosynthesis’ term is enriched in the earlier time-point subgroup Up_16h (Fig 2C). On the other hand, ‘lignin biosynthetic process’ is enriched at 48 HPT suggesting that lignin precursors are generated mainly at 16 HPT and then their polymerization occurs at a later stage after Sirodesmin PL application. This analysis additionally suggests that this toxin induces the upregulation of genes associated with indole glucosinolates metabolism (Supplementary Table S6).

Transcript suppression associated to Sirodesmin PL treatment was clearly enriched in photosynthesis-related GO terms, independently of the time-point analyzed (Fig. 2D and Supplementary Table S6). In particular, the term response to auxin was enriched at 48 HPT, which will be further analyzed in the next section.

Together these results suggest that Sirodesmin PL treatment leads to an induction of the *B. napus* immune response, and in particular against a fungal pathogen, while strongly suppressing the expression of photosynthesis-related genes. Further experiments shown below allow establishing if these transcript level differences translate into affecting those functions.

### Genes related to jasmonic acid and ethylene signaling are up-regulated in *B. napus* cotyledons in response to exposure to Sirodesmin PL

Plants rely on hormone signaling to activate their defense mechanisms against various stressors. Therefore, we investigated the expression of genes involved in the biosynthesis and signaling of plant hormones such as jasmonic acid (JA), ethylene (ET), salicylic acid (SA), gibberellic acid (GA), cytokinin (CK), abscisic acid (ABA) and auxin (Figure 3). In agreement with GO term enrichment analysis (Figure 2C) we observed a clear induction in the expression of genes involved in the biosynthesis and signaling of JA. Specifically, we observed up-regulation of *LIPOXYGENASE GENES* (*LOX2*, *3*, *4*, and *6*), *ALLENE OXIDE SYNTHASE* (*AOS*), *ALLENE-OXIDE CYCLASE* (*AOC2*, *3*, and *4*), and *OPDA REDUCTASE* (*OPR1*, *2*, and *3*), all involved in JA biosynthesis. Furthermore, key genes involved in subsequent JA signaling, such as *JASMONATE-ZIM DOMAIN* (*JAZ1*, *3*, *5*, *6*, *9*, *10*, and *12*) and encoding TIFY 10B transcription factor (*TIFY10B*), were up-regulated as well.

**Figure 3.**
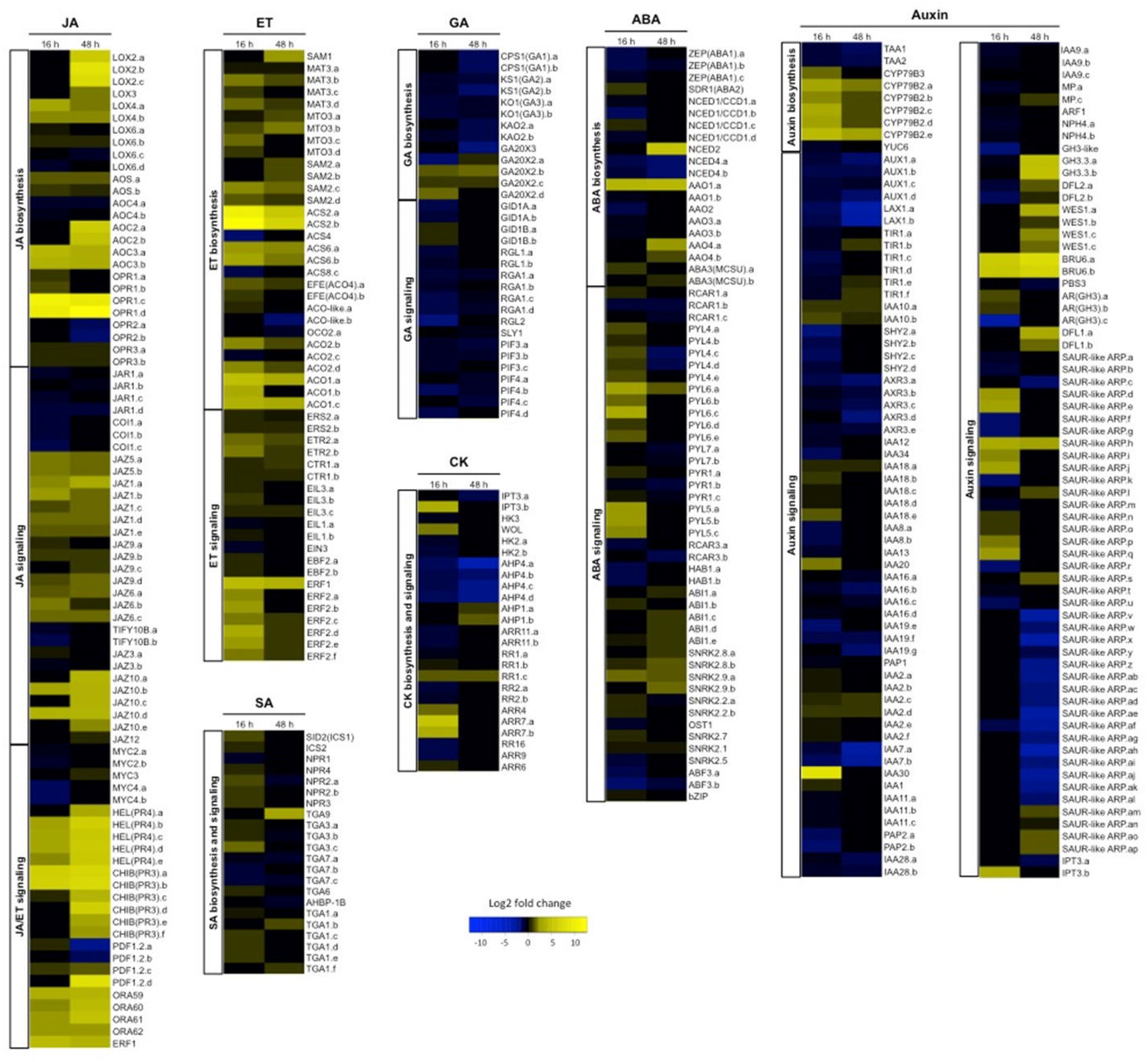
Expression of genes involved in the biosynthesis and signaling of plant hormones. Genes involved in the biosynthesis and signaling of jasmonic acid (JA), ethylene (ET), salicylic acid (SA), gibberellic acid (GA), cytokinin (CK), abscisic acid (ABA), and auxin were selected manually from the transcriptome analysis. Color bars ranging from blue to yellow represent downregulation and upregulation in transcript expression, respectively.

The expression of several genes involved in the biosynthesis of ethylene, another hormone involved directly in the plant defense mechanisms, such as *S-ADENOSYL-METHIONINE SYNTHETASE* (*SAM1* and *SAM2*), *METHIONINE ADENOSYLTRANSFERASE 3* (*MAT3*), *METHIONINE OVER-ACCUMULATOR 3* (*MTO3*), *1-AMINOCYCLOPROPANE-1-CARBOXYLIC ACID SYNTHASE* (*ACS2* and *ACS6*) and *1-AMINOCYCLOPROPANE-1-CARBOXYLIC ACID OXIDASE* (*ACO1, ACO2* and *ACO3*) were strongly up-regulated. In addition, genes involved in the signaling of the same hormone like *ETHYLENE RESPONSE SENSOR* (*ERS2*), *ETHYLENE RECEPTOR* (*ETR2*), *CONSTITUTIVE TRIPLE RESPONSE* (*CTR1*), *ETHYLENE INSENSITIVE 3-like* (*EIL1* and *3*), *ETHYLENE*-*INSENSITIVE 3-binding F-box* (*EBF2*) and, *ETHYLENE RESPONSE FACTOR* (*ERF1* and *2*) were also up-regulated. Some key genes involved in defense mechanisms shared by both JA and ethylene signaling pathways showed changes in their expression after Sirodesmin PL treatment. In particular, *bHLH transcription factor MYC 3* (*MYC3*), *HEVEIN-like protein* (*HEL*/*PR4*), *BASIC CHITINASE* (*CHI-B/PR3*), *PLANT DEFENSIN* (*PDF1.2a, b* and *c*), and *APETALA2/ETHYLENE RESPONSE FACTOR (AP2/ERF) domain transcription factor ORA59* (*ORA59*) were also up-regulated.

Auxin is a key hormone in the regulation of plant development, growth and is involved in the response to different biotic stresses. Here, we find an up-regulation of gene expression of some transcripts involved in the biosynthesis of this hormone. These genes belong to a family responsible for suppressing auxin signaling related to IAA conjugation machinery like *INDOLE-3-ACETIC ACID-AMIDO SYNTHETASE GH3* (*GH3.3, WES1*, *BRU6, DFL1* and *2,* and *SAUR-like auxin-responsive protein family*).

Contrastingly, the SA signaling associated genes did not show major changes in plants challenged with Sirodesmin PL. Moreover, apart from certain specific genes, treatment with Sirodesmin PL did not induce a comprehensive alteration in the expression of genes associated with GA, CK, and ABA-dependent pathways.

### Sirodesmin PL treatment induces ROS and callose accumulation in *B. napus* cotyledons

Reactive oxygen species (ROS) play a crucial role in plant defense against pathogen attacks. The plant employs a balance between ROS generation and scavenging systems upon pathogen attack to trigger an effective defense response. Total ROS accumulation was evident by dihydrofluorescein diacetate staining of cotyledons treated with 25 µM Sirodesmin PL at 24 HPT (Figure 4). Fluorescence intensity values were converted into a height map (2.5D) allowing the reconstruction of total ROS accumulation using 2.5D imagery of the treated zone. Total ROS accumulation showed a high extension in the Z-direction in cotyledons treated with Sirodesmin PL (Figure 4), while it was negligible in untreated cotyledons at the same time point.

**Figure 4.**
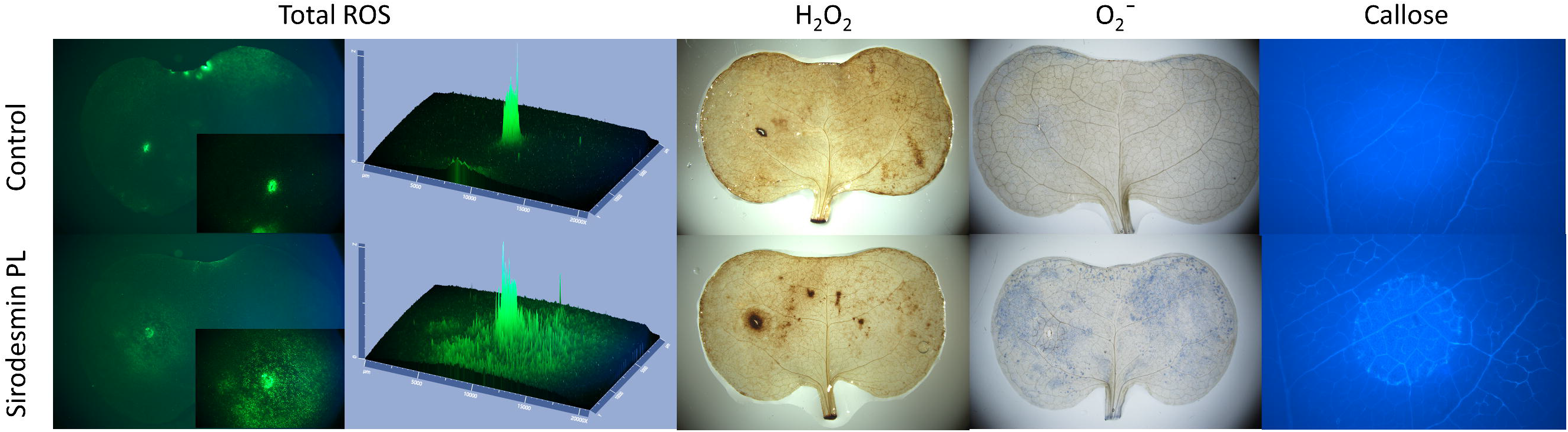
Sirodesmin PL-mediated induction of biochemical responses associated with the defense response. B. napus cotyledons were treated with 10 µl aliquots of a 25 µM Sirodesmin PL solution in ethanol/water (1:99). For the control condition, cotyledons were treated with 10 µl aliquots of a solution of ethanol/water (1:99). Fluorescence microscopy was used to analyze the accumulation of reactive oxygen species in cotyledons after 2’,7’-dichlorofluorescein diacetate (DCFDA) staining and callose deposition after aniline-blue staining. Light microscopy was used to analyze the accumulation of hydrogen peroxide and superoxide in cotyledons after 3’,3’-Diaminebezidine (DAB) and nitro blue tetrazolium (NBT) staining, respectively.

*B. napus* cotyledons exhibited a high increase in 3,3′-diaminobenzidine (DAB) staining after treatment with Sirodesmin PL, indicating a specific accumulation of hydrogen peroxide in the treatment area compared with untreated cotyledons (Figure 4). The production of superoxide radicals in plants treated with the toxin was evaluated by nitro blue tetrazolium (NBT) staining and evidenced by the precipitation of blue formazan within tissues. Cotyledon treatment with 25 µM Sirodesmin PL induced a significant increase in the level of superoxide radical at 24 HPT compared with untreated cotyledons (Figure 4). NBT staining reveals that the treatment induces a widespread and systemic accumulation of superoxide ions in a significant portion of the cotyledon.

Callose accumulates in the cell wall in response to plant pathogens (German *et al*., 2023). In order to test the alteration of callose deposition in *B. napus* plants treated with Sirodesmin PL, cotyledons were stained with aniline blue dye. Deposition of callose was evident at a short time point (16 HPT) in the site of Sirodesmin PL treatment (data not shown). At a later time point (48 HPT), an intense staining was observed in all the area of treatment with the toxin, not only in main cotyledons veins but also in the parenchymatic tissue (Figure 4).

All these experiments demonstrate that plant tissue responds rapidly to the toxin treatment leading to the generation of ROS and callose deposition.

### The expression of photosynthesis-related genes are down-regulated in cotyledons treated with Sirodesmin PL

The identification of an enrichment of GO terms related to photosynthesis among genes down-regulated as a consequence of Sirodesmin PL treatment (Figure 2D) prompted us to investigate in depth those genes involved in this process. From the list of DEGs initially identified, nuclear genes involved in the photosynthetic process were re-classified in Photosystem I and II (PSI and PSII), Intersystemic electron transport, Light-harvesting complex I and II (LHCI and LHCII) and ATPase complex. Expression of most of these genes was found to be down-regulated in cotyledons treated with Sirodesmin PL at both time points post-treatment (Figure 5). From a total of 179 photosynthesis-related genes, the expression of only 15 genes (8.4%) was upregulated. As shown in Figure 5, 13 of 39 genes belonging to the Intersystemic electron transport process were up-regulated in at least one time point. This means that with the exception of a few genes there is a global downregulation of genes related with this important biological process. Overall, this analysis indicates that Sirodesmin PL represses photosynthesis-related gene expression.

**Figure 5.**
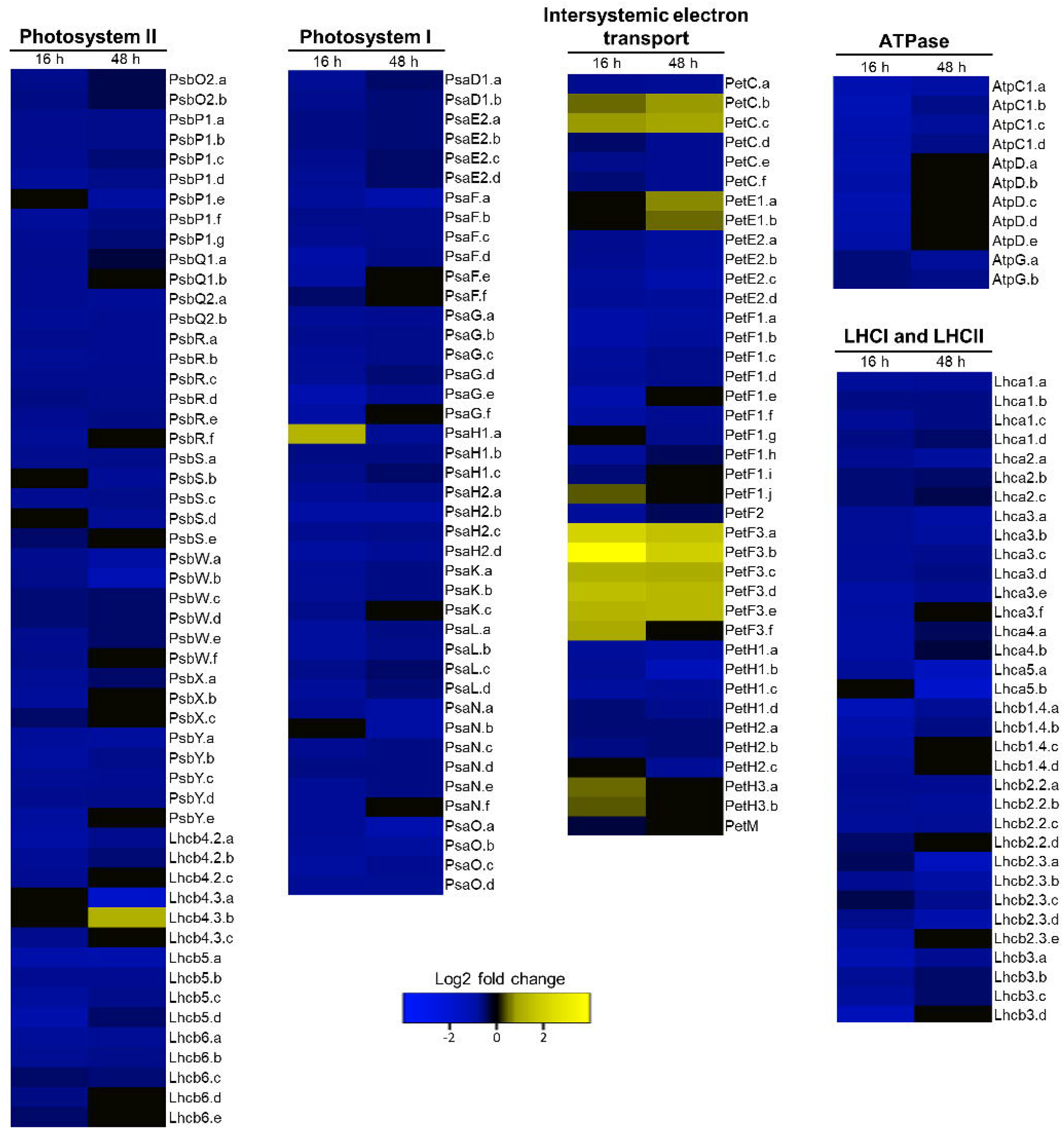
Expression of photosynthesis-related genes codified in the nucleus. Genes involved in the photosynthetic process were re-classified in Photosystem I and II (PSI and PSII), Intersystemic electron transport, Light-harvesting complex I and II (LHCI and LHCII, respectively) and ATPase complex were selected manually from the transcriptome analysis. Color bars ranging from blue to yellow represent downregulation and upregulation in transcript expression, respectively.

### Treatment with Sirodesmin PL negatively affects Photosystem II (PSII) activity

A decrease in photosynthesis-related transcripts associated to immunity activation may not necessarily coincide with a decrease in the abundance of corresponding proteins and consequently photosynthesis could still be unaffected (Huot *et al*., 2014). Taking this into account, we evaluated PSII activity at the physiological level by measuring chlorophyll transient fluorescence using an OJIP test. Fast chlorophyll *a* fluorescence rise kinetics OJIP-test analysis has been widely used to examine the structure, conformation and function of the photosynthetic apparatus (Chen *et al*., 2014). As shown in Figure 6A, the fluorescence rise transients obtained from control cotyledons at both time points showed a typical OJIP shape. Treatment with Sirodesmin PL induced changes in the OJIP curves in comparison to those observed in control samples (Figure 6C and D). The major change induced by Sirodesmin PL was a time-dependent increase of the J-step, which can be observed not only from OJIP curves but also as a result of the ratio between the different conditions analyzed, represented as black curves (Figure 6B and D). An increase in the fluorescence of the level of J-step is a result of the large accumulation of primary quinone electron acceptor (Q_A_) in PSII reaction centers (RCs), which is attributed to the interruption of the electron flow from the Q_A_ and secondary quinone electron acceptor (Q_B_).

**Figure 6.**
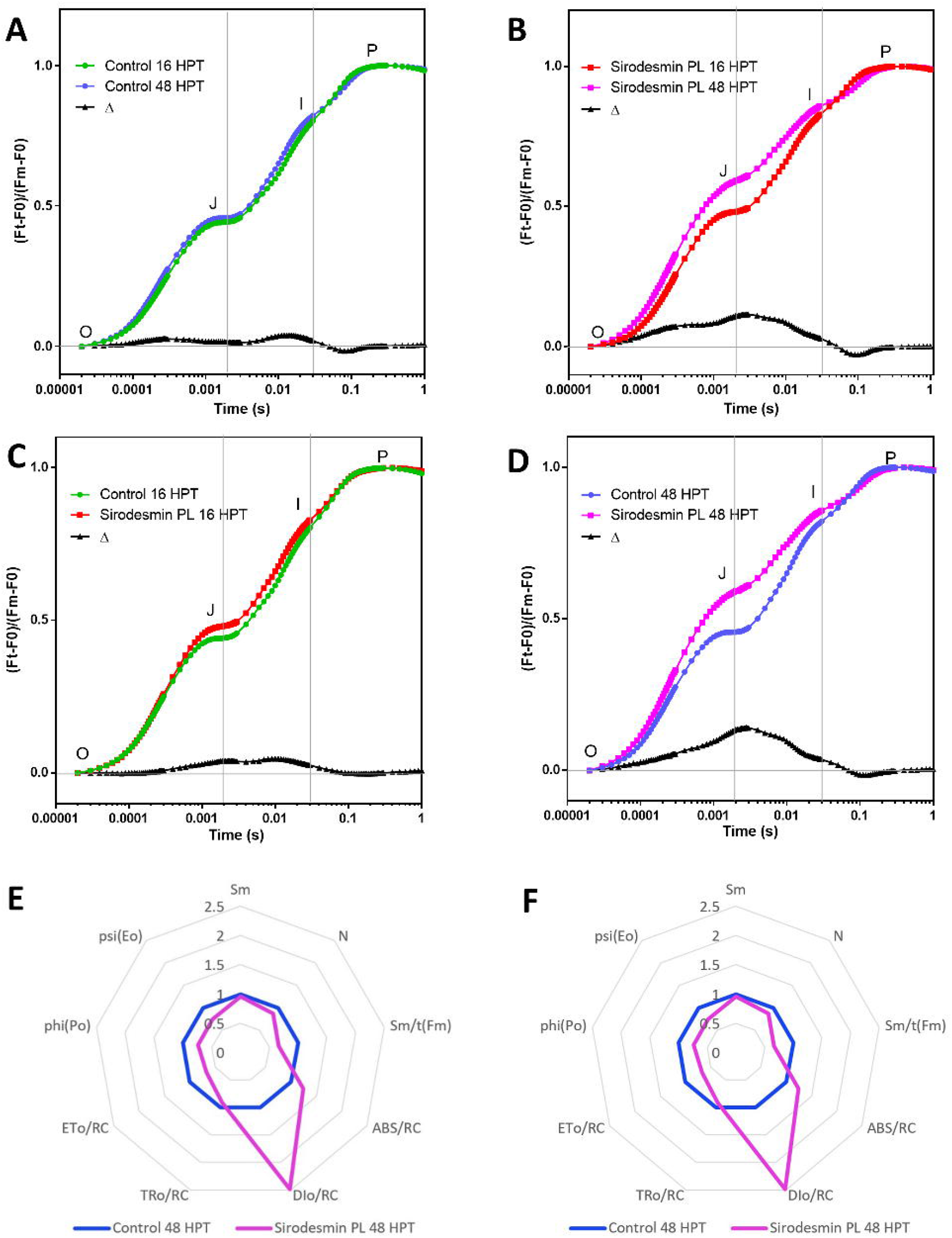
Chlorophyll *a* fluorescence transients of dark-adapted cotyledons of *B. napus* treated with Sirodesmin PL. The photosystem II activity was evaluated by measuring the Chlorophyll *a* fluorescence transients, analyzing the OJIP curve. *B. napus* cotyledons were treated with 10 µl aliquots of a 25 µM Sirodesmin PL solution in ethanol/water (1:99). For the control condition, + Measurements were performed at 16 and 48 hours after treatments. **A - D.** Raw fluorescence kinetics of cotyledons in vivo for control and treated samples at indicated times. The delta between different analyzed conditions is represented as a black curve. **A.** OJIP curve of control cotyledons at 16 (green) and 48 (blue) hours after treatment. **B.** OJIP curve of cotyledons treated with 25 µM Sirodesmin PL at 16 (red) and 48 (magenta) hours after treatment. **C.** OJIP curve of cotyledons control (green) and treated with 25 µM Sirodesmin PL (red) at 16 hours after treatment. **D.** OJIP curve of cotyledons control (blue) and treated with 25 µM Sirodesmin PL (magenta) at 48 hours after treatment. **E and F.** Spider graph presentation of some important OJIP-test parameters quantifying the behavior of PSII of cotyledons treated with 25 µM Sirodesmin PL (magenta) at 48 hours after treatment. All values are normalized to the values of the control (blue). A summary of the OJIP parameters used in this study is shown in the Supplementary Table S7.

To study the functionality of PSII and the transport of electrons between Q_A_ and Q_B_, we generated spider graphs with 48 HPT data from OJIP curves. As shown in Figure 6E and F, treatment with Sirodesmin PL negatively affects several processes such as the maximum quantum yield for PSII primary photochemistry (Phi (Po)), the probability that a trapped exciton moves an electron into the electron transport chain beyond Q_A_ (Psi (Eo)), the electron transport flux per reaction center and cross section (ETo/RC and ETo/CS, respectively), and the average fraction of open RCs of PSII in the time span between 0 to tFM (Sm/t(F_M_)). Finally, an increase in the dissipation of heat per excited CS and RC was observed.

### Sirodesmin PL treatment induces a decrease of the maximum quantum efficiency of PSII

Cotyledons treated with Sirodesmin PL showed a significant decrease of the maximum quantum efficiency of PSII at 48 HPT, as denoted by the variable fluorescence/maximum fluorescence ratio (Fv/Fm), which reflects damage to PSII (Figure 7A). No significant differences were observed at 16 HPT, but a significant decrease of 28% in the ratio Fv/Fm was observed as a consequence of the toxin treatment at the longer time point of 48 HPT.

**Figure 7.**
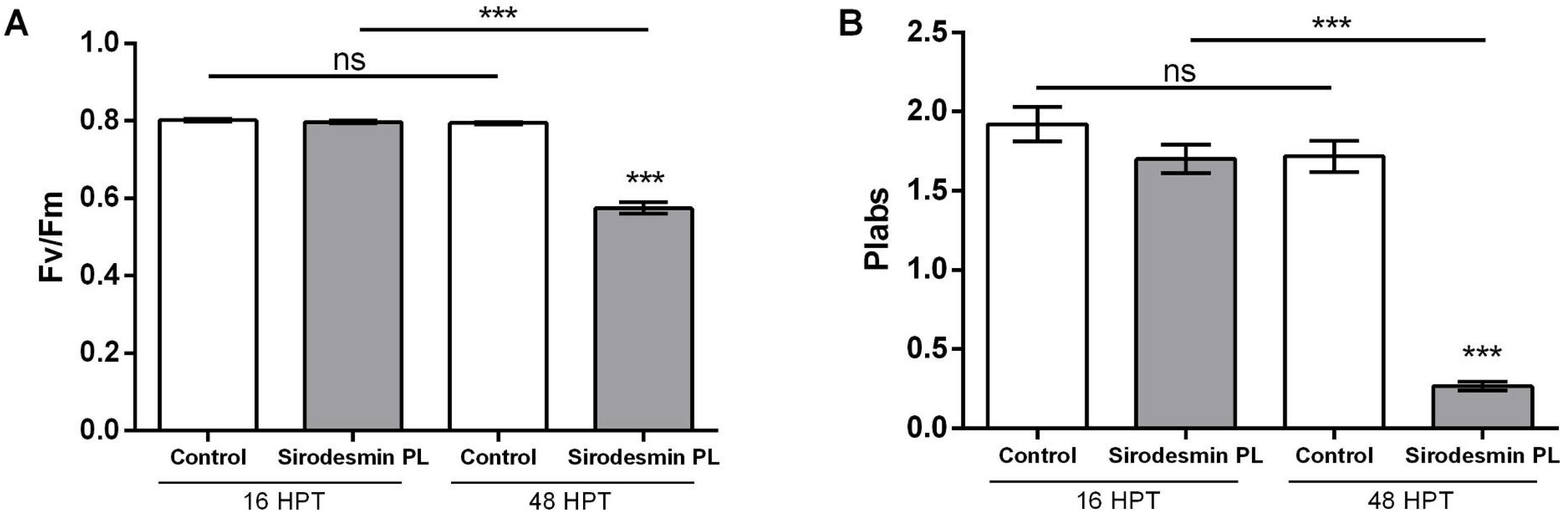
Analysis of Chlorophyll a fluorescence parameters. *B. napus* cotyledons were treated with 10 µl aliquots of a 25 µM Sirodesmin PL (grey bars) solution in ethanol/water (1:99). For the control condition (white bars), cotyledons were treated with 10 µl aliquots of a solution of ethanol/water (1:99). **A.** Fv/Fm, the maximum quantum yield of primary PSII photochemistry, was determined at the indicated times. **B.** PIabs, a performance index for energy conservation from absorbed photons to reduction QB, was determined at the indicated times. Results represent the mean of five to seven replicate cotyledons ± standard deviation, and statistically significant differences between Sirodesmin PL and the control condition, as determined by t-test, is indicated as *** (P ≤ 0.001). Upper lines shows statistically significant differences between Sirodesmin PL at 16 and 48 hours post treatment and the control condition at 16 and 48 hours post treatment, as determined by t-test, is indicated as *** (P ≤ 0.001).

The most sensitive OJIP-test parameter that responded to the treatment of Sirodesmin PL was the performance index (potential, PIabs) for energy conservation from photons absorbed by PSII to the reduction of intersystem electron acceptors. In concordance with Fv/Fm, a drastic decrease of 84.3% in the PIabs was detected after 48 HPT (Figure 7B). No significant differences were observed between treatments at short time of exposure. The PIabs is regulated by three functional steps, which are absorption of light energy (AB), trapping of excitation energy (TR) and conversion of excitation energy (ET). Thus PIabs is a product of the three independent components: PI_ABS_= γ_RC_/(1-γ_RC_) · ϕ_Po_/(1-ϕ_Po_) · ψ_Eo_/(1-ψ_Eo_). Here, we observed significant changes in the three components at 48 HPT (Supplementary Fig. S2).

Collectively these results indicate that the transcriptional down regulation of photosynthesis-related genes correlate with a severe disruption of photosynthesis, particularly at 48 HPT.

### The number of chloroplasts per cell decrease in cotyledons treated with Sirodesmin PL

Taking into account the results showed in previous sections, we decided to evaluate by microscopy the presence of morpho-cytological changes using cotyledon semiithin sections. In untreated cotyledons a mean of 18 and 13 chloroplasts per cell were found in the spongy parenchyma and palisade cell layer, respectively. At 48 HPT the number of chloroplasts per cell on cross-sections of the spongy parenchyma decreased significantly (27.3%) in cotyledons treated with Sirodesmin PL compared to control samples. A similar trend was observed on the palisade cell layer, in that 25.6% less chloroplasts were detected on cross-sections of cotyledons exposed to Sirodesmin PL (Figure 8).

**Figure 8.**
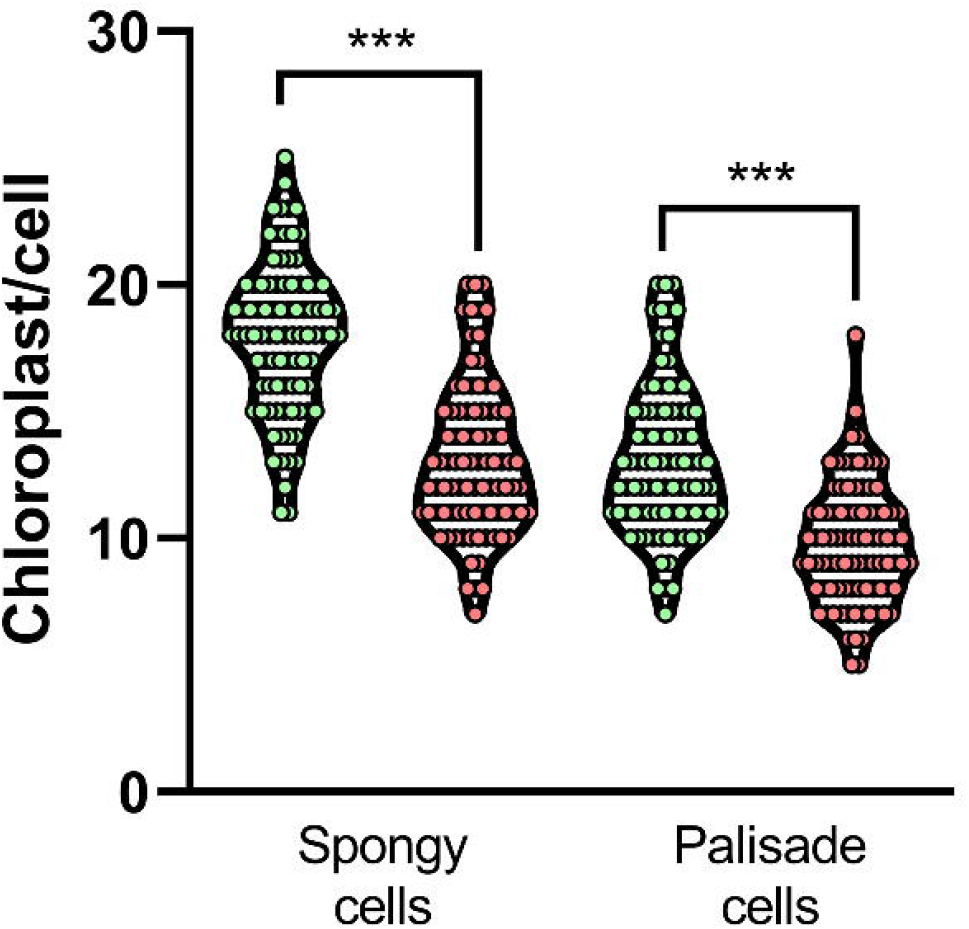
Changes in the relative number of chloroplasts. The number of chloroplasts was evaluated per cell on longitudinal semithin-sections of the palisade cell layer and the spongy parenchyma in *B. napus* cotyledons treated with 25 µM Sirodesmin PL (red dots) and compared to the control (green dots). Measurements were performed at 48 hours post-treatment. Violin graph show the distribution of the number of chloroplasts per cell and the means ± standard deviation. Significant differences between Sirodesmin PL treated and control samples, as determined by t-test, are indicated as *** (P ≤ 0.001).

## Discussion

Phoma Stem Canker is one of the most significant diseases to impact agriculture worldwide, affecting oilseed rape cultivation and causing severe economic losses (Fitt *et al*., 2006). This disease is primarily caused by the hemibiotrophic fungus *L. maculans*, which employs both biotrophic and necrotrophic stages during its complex infective cycle (Howlett *et al*., 2001). During this interaction, if the plant possesses a resistance (R) protein that is able to recognize the corresponding pathogen avirulence (Avr) gene product, an incompatible interaction occurs that arrests the pathogen (Becker *et al*., 2019). Like most phytopathogenic fungi, *L. maculans* utilizes various secondary metabolites, such as phytotoxins, that contribute to virulence and play distinct roles throughout the infection process (Pedras *et al*., 1990; Pedras and Biesenthal, 1998). Simultaneously with the global boom in oilseed rape production in the 1990s, numerous studies assessed the negative impact of Phoma stem canker on crop productivity. During those years, the scientific community classified *L. maculans* as Tox+ (highly virulent/Group A) or Tox0 (weakly virulent/Group B) based on its ability to produce Sirodesmin PL (reviewed in (Rouxel *et al*., 1994)). Tox0 was reclassified as *L. biglobosa* (Shoemaker and Brun, 2001). These studies indirectly highlighted the significance of Sirodesmin PL in the virulence of *L. maculans*. Although the genetics of the *L. maculans–B. napus* interaction has been vastly studied, the role of Sirodesmin PL is largely unresolved.

Here, we demonstrate through trypan blue staining that treatment with Sirodesmin PL induces cell death in *B. napus* cotyledons. This effect is concentration-dependent and time-post application-dependent (Figure 1 and Supplementary Figure S1). A similar effect was previously reported in one of the early studies describing the toxicity of this toxin in *B. napus* and other hosts of *L. maculans* (Badawy and Hoppe, 1989). However, in our system, the toxic effect could be observed at considerably lower concentrations than previously reported, which may be related to differences in toxin purification and quantification methods.

Some evidence suggests that Sirodesmin PL and other compounds belonging to the ETP group of fungal natural products are capable of inducing cell damage due to the high reactivity of the disulfide bridge, which has the ability to interact with cysteine residues present in proteins, thus affecting their proper function (Gardiner *et al*., 2005*a*). Gliotoxin is one of the most studied ETP toxins, produced by the opportunistic pathogenic fungus *Aspergillus fumigatus*. The role of the disulfide bridge in the mechanism of action of this toxin has been extensively analyzed (Scharf *et al*., 2016). The gene cluster involved in gliotoxin biosynthesis exhibits a high similarity to that found in *L. maculans* for the synthesis of Sirodesmin PL, demonstrating overlap in the overall biosynthesis mechanisms of both compounds (Gardiner *et al*., 2005*a*). Here, we show that the permanent reduction of the disulfide bridge of Sirodesmin PL, using BH_4_Na, completely eliminates the damage caused by the toxin (Figure 1). In line with these results, some evidence indicates that the presence of the oxidized disulfide bridge is crucial for gliotoxin entry into cells (Schrettl *et al*., 2010). Furthermore, recently Hou et al. (2023) showed that the reduction of the disulfide bridge of Deoxyverticillin A, another ETP metabolite produced by *Clonostachys rogersoniana*, fails to induce apoptosis and genomic instability in HeLa cells. These findings suggest that the disulfide bridge is essential for toxins belonging to the ETP group to enter the host cells and to react with proteins. Our results indicate that Sirodesmin PL follows this rule although additional assays are necessary to confirm it.

No studies have reported global transcriptional modifications in *B. napus* plants treated with Sirodesmin PL. Interestingly, the results obtained in this study reveal changes in the expression levels of thousands of genes in *B. napus* cotyledons at both short and long times after treatment with Sirodesmin PL (Figures 2A-B). Classification of the induced DEGs into GO terms showed a significant enrichment at both time points of up-regulated genes associated with different processes related to plant defense. Among them are ‘Response to chitin’, ‘Sulfur compound biosynthetic process’, ‘Toxin metabolic processes’, ‘Secondary metabolic and catabolic processes’, ‘Immune system process’, and ‘Response to oxidative stress’ (Figure 2C). In this context, an up-regulation of genes encoding transcription factors was observed at 16 and 48 HPT (also shared at both times). Genes encoding Zinc finger transcription factors, Ethylene-responsive element binding proteins, and WRKY proteins, among others, were up-regulated by treatment with Sirodesmin PL. Additionally, the up-regulation of genes associated with biological processes encode proteins participating in the ubiquitination pathway, such as U-BOX and Toxicos en Levadura, while the down-regulation of genes associated with photosynthetic processes (Supplementary table S5 and S6) resembles typical responses to chitin treatment (Zhang *et al*., 2002; Libault *et al*., 2007). Further studies are necessary to elucidate a possible relationship between Sirodesmin PL recognition and the activation of PAMP-triggered immunity. The ‘Sulfur compound biosynthetic process’ was up-regulated at both post-treatment times (Figure 2C, Supplementary table S5). Several genes associated with this process encode enzymes involved in glutathione metabolism (glutathione-synthase, -reductase, and -transferase). High intracellular levels of glutathione are necessary to maintain a correct redox balance but also to reduce the disulfide bridge of ETP toxins and intensify gliotoxin-induced cytotoxicity (Bernardo *et al*., 2003; Carberry *et al*., 2012). As shown in Figure 1C, Sirodesmin PL enters the cell in an oxidized manner (S-S) and would then be reduced (-SH HS-) by the action of reduced glutathione (GSH), similar to what happens with gliotoxin (Carberry *et al*., 2012), as a necessary step to induce cellular damage. The up-regulation of genes involved in glutathione metabolism could increase the intracellular levels of the reduced species. This increase could be part of an antioxidant response of the plant, providing an advantageous condition for *L. maculans* by reducing Sirodesmin PL during its infective process.

Upon recognizing a pathogen, plants activate various defense mechanisms to counteract its spread. Programmed cell death (PCD) constitutes a highly effective strategy to prevent the growth and dissemination of the pathogen throughout the rest of the plant. PCD is very effective against biotrophic and hemibiotrophic pathogens but not against necrotrophic ones that in some cases may trigger PCD as part of causing damage to the plant (Glazebrook, 2005). Our results indicate that Sirodesmin PL not only induces cell death and changes at the transcriptional level of genes involved in process related to response to oxidative stress (Supplemental tables S5 and S6), but also triggers the accumulation of reactive oxygen species, particularly H_2_O_2_ and superoxide ions (Figure 4). We have previously shown a similar response mechanism in Arabidopsis plants treated with Botrydial, a sesquiterpenoid toxin produced by *B. cinerea* (Rossi *et al*., 2011). This defense strategy extends to other toxins, including Victorin (*Cochliobolus victoriae*), PC-toxin (*Periconia circinata*), SnTox1 (*Parastagonospora nodorum*), ToxA (*Pyrenophora tritici-repentis*), BsToxA (*Bipolaris sorokiniana*), among others (Faris and Friesen, 2020). Another compound involved in defense mechanisms is callose, a structural polysaccharide located in cell walls that can accumulate to form a physical defense barrier against pathogens. Marked callose accumulation has been demonstrated in *B. napus* infected with *L. maculans*, indicating that the accumulation of this compound plays a role in defense (Liu *et al*., 2018). Here, we show that Sirodesmin PL induces the accumulation of callose (Figure 4), suggesting that the plant recognizes the toxin as a potential damaging agent, likely associated with pathogens. A similar response was reported by Liu et al. (2018), who demonstrated that callose accumulation occurs in *B. napus* cotyledons infected with *L. maculans* in both compatible and incompatible interactions. Additionally, several genes involved in callose synthesis were induced at both short and long times post-treatment with Sirodesmin PL (Supplementary table S3 and S4). The accumulation of various defense compounds (ROS and callose) suggests that complex signaling occurs in *B. napus* after recognizing Sirodesmin PL and correlates with the expression levels of genes involved in the production of these types of compounds (Supplementary table S3 and S4).

A balance in the levels of hormones involved in defense mechanisms is crucial for the effectiveness of plant defenses against microbes. Here, we provide evidence of a significant modification in the expression levels of genes involved in the synthesis and signaling of various hormones in *B. napus* cotyledons treated with Sirodesmin PL. In particular, a pronounced induction of genes associated with the hormones JA and ET was observed (Figure 3). These hormones are typical defense markers against necrotrophic microorganisms, such as *B. cinerea* or *S. sclerotiorum* (Glazebrook, 2005). Transcriptomic analysis of cotyledons infected with *L. maculans* demonstrates the induction of SA signaling during the biotrophic stage and the activation of JA signaling during the necrotrophic stage in this interaction (Haddadi *et al*., 2019; Borhan *et al*., 2022). Recently, Zaid et al. (2022) demonstrated that gliotoxin induces the transcription of genes associated with the same hormonal pathways, indicating that ETP toxins may trigger typical defense responses against necrotrophic pathogens (Padmathilake and Fernando, 2022). Several genes involved in auxin-mediated signaling were also induced after treatment with Sirodesmin PL. The direct involvement of auxins in defense processes is debated in the literature, although it is generally assumed that this hormone positively contributes to resistance against necrotrophic pathogens (Llorente *et al*., 2008). Interestingly, the transcription of the *INDOLE-3-ACETIC ACID-AMIDO SYNTHETASE GH3* gene family was induced. Some of these genes participate in the synthesis of other antimicrobial compounds such as camalexin and callose (Fu and Wang, 2011). Our results indicate that, although the expression of genes involved in the synthesis of auxin is up-regulated (Figure 3), the signaling process of this hormone is down-regulated (Figure 2D). In this way we cannot rule out that Sirodesmin PL represses auxin signaling and the plant tries to compensate for this repression by increasing the synthesis of the hormone. Thus, it appears that the observed cell death after the treatment with Sirodesmin PL is part of PCD processes, characterized by the accumulation of reactive oxygen species, callose, and the induction of JA- and ethylene-dependent mechanisms, as a result of typical defense against necrotrophic pathogens. In general, these responses closely resemble those found in incompatible and/or later (necrotrophic) stage of the interactions between *L. maculans* and *B. napus* (Sašek *et al*., 2012; Liu *et al*., 2018; Becker *et al*., 2019; Haddadi *et al*., 2019; Borhan *et al*., 2022).

Photosynthesis plays an indispensable role in energy production and this process undergoes various alterations under stressful conditions. Additionally, the chloroplast is the compartment where various hormones involved in defense processes are synthesized (Lu and Yao, 2018). Most plant-pathogen interactions tend to suppress photosynthesis, generally associated with overall malfunction, ultimately leading to PCD. Here, we demonstrate that approximately 90% of nuclear genes encoding proteins targeted to the chloroplast exhibit strong down-regulation, and this response was generalized in all components of the photosynthetic apparatus (Figure 5). A similar response was observed in the transcriptome of incompatible interactions between *B. napus* with *L. maculans* (Becker *et al*., 2019). Considering the results observed in our transcriptome analysis and the down-regulation of numerous genes associated with different photosynthetic processes, we decided to evaluate this process from a functional perspective. As shown in Figure 6, treatment with Sirodesmin PL leads to a significant decrease in transient Chl *a* fluorescence, indicating malfunction of PSII. Similarly, Tenuazonic acid (TeA), a phytotoxin produced by the fungus *Alternaria alternata*, strongly inhibits PSII activity (Chen *et al*., 2014). This toxin, like Sirodesmin PL, causes a marked increase in the J-step of the typical OJIP curve, which corresponds to peak concentrations of Q_A_^-^QB and Q_A_^-^Q_B_^-^ formed by electron transport from Q_A_ to Q_B_ (Stirbet and Govindjee, 2011). This observation is in concordance to the down-regulation of genes associated with the PSII (Figure 5). Dissection of PSII fluorescence data indicated that *B. napus* cotyledons treated with Sirodesmin PL contained fewer active reaction centers (RCs) per leaf cross-section (CS) than untreated cotyledons (Figure 6F), suggesting that fewer pigments were associated, at least in part, with photosynthetic complexes. These results are in the same line with those obtained from transcriptome analysis, as GO term ‘Pigment biosynthetic process’ was enriched among down-regulated DEGs at 16 HPT (Figure 2D). The lower RC/CS ratios of treated cotyledons resulted in the distribution of incident light through fewer RCs, leading to higher photons being absorbed (ABS/RC) and higher energy being dissipated (DIo/RC) per active RC. Dissipated energy can manifest in two ways: through photochemical and nonphotochemical processes. The latter can lead to cellular damage via energy transfer that excites chlorophyll and cellular components such as phospholipids and proteins. Complementary, PI_ABS_ and Fv/Fm values are fluorescence parameters that provide complete and quantitative information about vitality of plants (Strasser and Stirbet, 2001). The drop in those parameters following Sirodesmin PL treatment (Figure 7) indicates that this toxin has a negative impact on energy trapping and electron flow. It also blocks the Qb site, thereby contributing to the adverse effects of this stress condition on the photosynthetic machinery. In rice plants subjected to salt stress, the decrease in PIabs parameters was found to be highly correlated with an increase in cellular damage, as indicated by higher levels of malondialdehyde content measured using the TBARs method (Rachoski *et al*., 2015). These parameters drop after Sirodesmin PL treatment (Figure 7) indicating that this toxin affects negatively the photosynthetic electron flow, contributing thus to the adverse effects of this stress condition on the photosynthetic machinery.

Previously, we determined a similar pattern in changes in photosynthetic capacity in tobacco plants infected with the necrotrophic fungus *B. cinerea* (Rossi *et al*., 2017). Pecan plants infected with the hemibiotrophic fungal species in the genus *Phomopsis* also show a decrease in chlorophyll fluorescence (Mantz *et al*., 2021). It seems that the decline in photosynthetic activity is the result of a multifaceted process involving a decrease in transcript levels of genes encoding proteins involved in photosynthesis and also a decrease in the number of chloroplasts per cell, both in spongy parenchyma and palisade parenchyma (Figure 8). A similar response has been reported in rice plants infected with the necrotrophic fungus *Rhizoctonia solani*. In this pathosystem, infection by this pathogen causes significant alterations at the chloroplast level, accompanied by a decrease in the expression of genes associated with photosynthesis and parameters associated with this activity (Ghosh *et al*., 2017). Additionally, several phytotoxins produced by *Pseudomonas syringae* target chloroplasts, inducing structural and functional alterations and chlorosis at the tissue level (Bender *et al*., 1999), similar to our results with Sirodesmin PL. In this way, the drop in photosynthetic activity, the increase in dissipated energy and the decrease in the number of chloroplasts per cell could increase ROS to lethal levels causing the cell death observed after treatment with Sirodesmin PL (Simon *et al*., 2013; Zechmann, 2019).

In conclusion, our results indicate that Sirodesmin PL is an important factor used by *L. maculans* in its host-pathogen interaction, capable of causing cell death as the final step in its mechanism of action. In its native (oxidized) form, this toxin enters to the *B. napus* cells, which attempt to activate defense mechanisms characterized mostly by the generation of ROS and callose deposition, which is accompanied by significant transcriptional modifications. However, as part of the ongoing battle between plants and pathogens, *L. maculans* would use Sirodesmin PL to induce cell death for its own benefit, necessary to complete the necrotrophic cycle. Cell death could be a consequence of the down-regulation of genes associated with photosynthesis and energy generation, resulting from the decrease in photosynthetic activity as well as the number of chloroplasts per cell. The accumulation of ROS, combined with the processes mentioned above, would lead to a typical PCD favoring the infective process of *L. maculans*.

## Supplementary data

**Supplementary Figure S1.** Cell death in *B. napus* cotyledons induced by Sirodesmin PL at 16 and 48 HPT.

**Supplementary Figure S2.** Statistical analysis of Performance Index (PIabs) components. A-absorption of light energy (AB), B-trapping of excitation energy (TR) and C-conversion of excitation energy (ET).

**Supplementary Table S1.** Summary of sequencing data for each of the libraries generated in this work.

**Supplementary Table S2.** Pearson correlation analysis comparing all the sequenced samples

**Supplementary Table S3.** Genes expressed at 16 HPT in at least one condition (RPKM ≥ 2)

**Supplementary Table S4.** Genes up-regulated by sirodesmin PL toxin at 48 HPT (fold change ≥2; q < 0.05)

**Supplementary Table S5.** Gene expression values of up-regulated, down-regulated and shared at 16-and 48 HPT.

**Supplementary Table S6.** Gene Ontology (GO) enrichment analysis.

**Supplementary Table S7.** Summary of the OJIP test formulae

## Acknowledgements

MAP, HGR, SM, FMR, AG, OAR and FRR are members of the Research Career of CONICET, Argentina. SM is a doctoral fellow of FONCyT, Argentina. EC and IA are members of School of BioSciences, University of Melbourne, Australia. The authors are grateful to Beatriz Wiss (CPA-CONICET, Argentina) and Juan Ezquiaga (CIC, Buenos Aires province, Argentina) for valuable technical assistance.

## Funding

This work was financially supported by grants of Agencia Nacional de Promoción Científica y Tecnológica (PICT Startup 2020-0053 and PICT 2019-1137) and CONICET (PIBAA 28720210100989CO) to FRR

## Notes

### Competing Interest Statement

The authors have declared no competing interest.

